# A Personalized Switched Systems Approach for the Optimal Control of Ventricular Assist Devices based on Atrioventricular Plane Displacement

**DOI:** 10.1101/2020.05.27.119149

**Authors:** Clemens Zeile, Thomas Rauwolf, Alexander Schmeisser, Jeremi Kaj Mizerski, Rüdiger C. Braun-Dullaeus, Sebastian Sager

## Abstract

**Objective:** A promising treatment for congestive heart failure is the implementation of a left ventricular assist device (LVAD) that works as a mechanical pump. Modern LVADs work with adjustable constant rotor speed and provide therefore continuous blood flow; however, recently undertaken efforts try to mimic pulsatile blood flow by oscillating the pump speed. This work proposes an algorithmic framework to construct and evaluate optimal pump speed policies.

**Methods:** We use a model that captures the atrioventricular plane displacement, which is a physiological indicator for heart failure. We employ mathematical optimization to adapt this model to patient specific data and to find optimal pump speed policies with respect to ventricular unloading and aortic valve opening. To this end, we reformulate the cardiovascular dynamics into a switched system and thereby reduce nonlinearities. We consider system switches that stem from varying the constant pump speed and that are state dependent such as valve opening or closing.

**Results:** As a proof of concept study, we personalize the model to a selected patient with respect to ventricular pressure. The model fitting results in a root-mean-square deviation of about 6 mmHg. Optimized constant and piecewise constant rotor speed profiles improve the default initialized solution by 31% and 68% respectively.

**Conclusion:** These in silico findings demonstrate the potential of personalized hemodynamical optimization for the LVAD therapy.

**Significance:** LVADs and their optimal configuration are active research fields. Mathematical optimization enhances our understanding of how LVADs should provide pulsatility.

## I. Introduction

LEFT VENTRICULAR ASSIST DEVICES (LVADs) provide mechanical circulatory blood support and have become a well-established and successful therapy for end-stage heart failure patients with estimated more than 5000 implanted pumps annually worldwide [1], [2]. The role of the heart assist devices is growing in recent years since there are major improvements in the long term treatment [3]. Contemporary LVADs implement rotary continuous blood flow and are internally implanted in contrast to pulsatile and extracorporeal pumps, which represent the original LVAD design, but which are bigger, less durable, and more invasive than their continuous flow counterpart [4].

These either axial- or centrifugal-flow pumps were originally designed to apply a fixed constant rotary speed. However, there is evidence that this lack of pulsatile flow can cause numerous adverse effects that include gastrointestinal bleeding [5], reduced end-organ function [6], aortic valve thrombosis and de novo aortic insufficiency [7]. To this end, the latest generation of devices features a pulsatile mode in addition to the constant speed option that oscillates the motor rotation speed periodically for a short period of time before returning to the constant speed operation. Examples of these devices and modes are HeartMate 3 with the *Pulse mode,* HeartWare HVAD with the *Lavare Cycle* and EXCOR/INCOR [8]. For further details on the devices, the medical background, therapy planning and prognosis we refer to the reviews [9]–[12].

A vast amount of preclinical models for the evaluation and testing of LVADs via pump speed modulation has been proposed. Amacher et al. [13] reviewed a range of studies [14]–[17] where a preselected constant, sine or square wave speed profile is assumed. Chosen parameters were adjusted for amplitude and phase shift in order to analyze the effect on relevant physiological quantities. Specifically, high speed pumping during ventricular contraction, also denoted as copulsative mode, were found to be beneficial in terms of pulsatility in the systemic arterial circulation. Counterpulsative pumping, i.e., low speed pumping during the ventricular contraction, enhanced left ventricle (LV) unloading [18].

A preselected speed profile does not adjust to dynamic changes in the state of the cardiovascular system. For this reason, control strategies for the blood pumps were developed that take into account different physiological objectives and which were classified in the review of Bozkurt [19]. Physiological control following the Frank-Starling mechanism by pumping preload dependent has been proposed in [20]–[22]. Control algorithms that aim for unloading the LV were elaborated in [23], [24]. Speed regulation algorithms for generating sufficient perfusion and detecting ventricular suction [25], [26] or pulmonary oxygen gas exchange tracking [27] are other goals and, finally, multi-objective variants exist [28].

Due to the increased necessity of LVADs for clinical use, a wide range of different methods from control engineering has been applied such as adaptive [29], [30], robust [31], model predictive [32], fuzzy logic [33], proportional integral derivative [34], sliding mode [35] and iterative learning control [36]. We refer to [37] for a detailed review.

In this work, we follow an optimal control approach since it offers a flexible framework to include and combine multiple objective and constraint functions. So far, optimal control studies based on cardiovascular system modeling appear to be very limited in the context of ventricular assist devices. [38] investigated the use of LVADs for preload manipulation maneuvers in animal trials. We build on [39], where the continuous pump speed profile is found with an optimal control algorithm based on a lumped cardiovascular system model and compared with both a constant and a sinusoidal-speed profile. In contrast, we do only numerical simulation and no verification with a mock circulation system. Our idea is to consider the cardiovascular dynamics as a system that *switches* between different phases in a single cardiac cycle, e.g. by valve opening or closing, in each of which different dynamics apply. We use solving techniques tailored for *switched systems* to reduce the underlying system nonlinearities and leverage the computations. Within this framework, we present a novel algorithm to calculate optimal piecewise constant (pwc) pump speed modulation in accordance with the above mentioned pulsatility modes for modern devices and with respect to ventricular unloading and opening of the aortic valve. For comparison, we compute the optimal continuous and constant speed profiles. Furthermore, we consider to adapt model parameters to patient specific data with a nonlinear regression objective function so that we deal with a personalized model.

Another fundamental difference of our approach to [39] and all other model based approaches lies in the used model: Instead of applying a time-varying elastance function to represent the pressure-volume (PV) relationship in heart chambers, we base our model on the contribution of the longitudinal atrioventricular plane displacement (AVPD) to ventricular pumping, which is novel in the LVAD context. It has been established that the atrioventricular (AV) plane behaves as a piston unit by moving back and forth in the base-apex direction, creating reciprocal volume changes between atria and ventricles [40]. Also, there is strong evidence that the magnitude of AVPD is a reliable index for diagnosis of heart failure [41]. Since elastance functions cannot explain the behavior of ventricular walls and fail to simulate the interaction between the LV and an assist device [42], [43], we reuse and extend an AVPD model introduced in [44] and altered to the switched systems setting in [45]. Alternatives for replacing the elastance model are myofiber or sarcomere mechanics approaches [46] as in the CircAdapt model [47], [48], though a great number of discontinuities and nonlinear equations limit their applicability to (gradient based) optimization and control techniques. The presented approach is clinically applicable, since the atrioventricular plane (AVP) motion is relatively easy to measure via noninvasive echocardiography.

The outline of this article is the following: We describe the cardiovascular and LVAD system model in Section II-A, before we define constraints in Section II-B, the clinical data and model personalization in Section II-C. Afterwards, we formulate the optimal control problem (OCP) in Section II-D and we define an algorithmic approach to solve it in Section II-E. We present simulation results in Section III and discuss the realistic and algorithmic setting together with limitations in Section IV. We wrap up the article with conclusions in Section V.

## II. Methods

### A. Cardiovascular System and LVAD modeling

This study uses a lumped model of the cardiovascular system based on the representation of the left heart. We combine the AVPD model as proposed and validated in [45] with an axial pump LVAD model that has been validated in [49]. The proposed model consists of nine ordinary differential equations (ODEs) for the pressure *P*(*t*) of left atrium (LA), LV, Aorta (A), systemic artery (S), and venous system (V), the flow *Q*(*t*) in the Aorta (A) and in the LVAD as well as the velocity *v*(*t*) of the AVPD and its position *s*(*t*), where (*t*) denotes the time dependency. The cardiovascular system can be steered with the continuous control *u*(*t*) that represents the rotary pump speed. The ODE system reads for *t* ∈ [*t*_0_,*t_f_*] ⊂ ℝ:

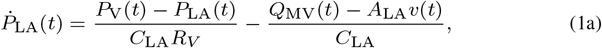

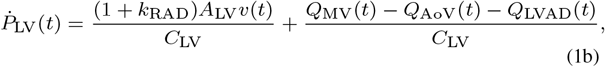

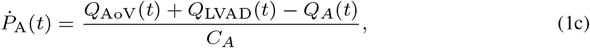

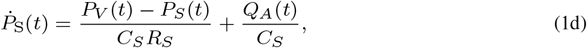

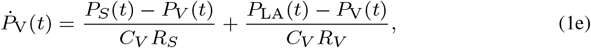

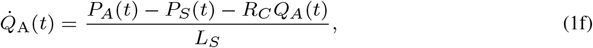

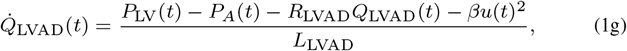

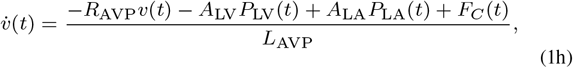

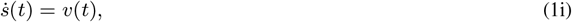

where the default parameter values for the compliances *C*, resistances *R*, and inertances *L* are given in the Supplemental Material I. The model uses the valve flows^1^ defined by

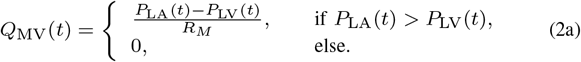

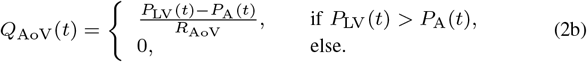

The AV plane contraction force is assumed to be a pwc function in the following sense

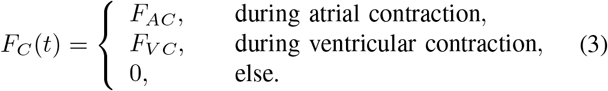

We specify in Section II-E how these contraction phases are mathematically defined and skip their formal introduction here.

Figure 1 gives a schematic overview of the lumped model of the heart and the circulatory system. In the following, we group the differential states into the vector x = [*P*_LA_, *P*_LV_, *P*_A_, *P*_S_,*P*_V_, *Q*_A_, *Q*_LVAD_, *v, s*]^T^ and write the dynamical system (1a)–(1i) as

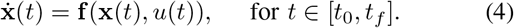

**Fig. 1.**
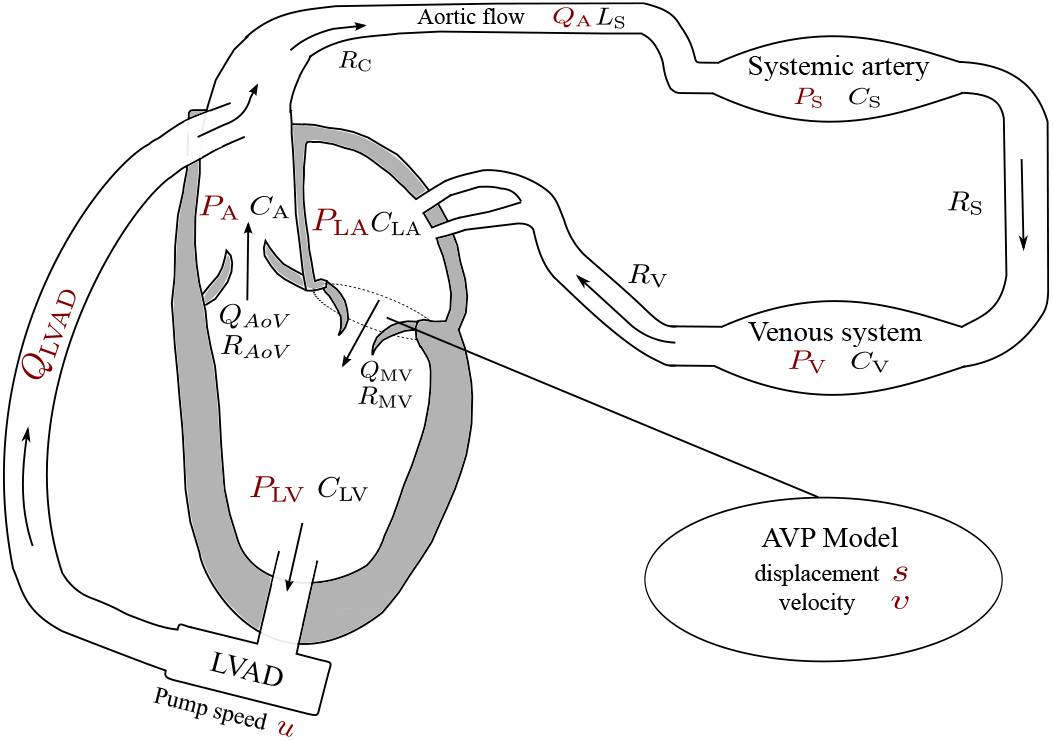
Illustration of the simplified model of the left heart, the circulatory system, and the LVAD. Differential states and the pump speed control *u*(*t*) are depicted in red. The cyclic flow is indicated by the arrows. The model consists of five compartments for the left atrium (LA), left ventricle (LV), Aorta (A), systemic artery, and venous system, represented with the pressure functions *P*(*t*). These variables interact with the flows *Q*(*t*) in the LVAD and the Aorta, while the AV interaction is modeled by the velocity *v*(*t*) and position *s*(*t*) of the atrioventricular plane displacement (AVPD). Compliance, resistance and inertance parameters *C, R, L* are depicted next to the corresponding compartment.

Figure 2 illustrates the AVPD model, where the AVP refers to the separating tissue between LV and LA that surrounds the mitral valve. During atrial contraction, the force *F_C_* pulls the AVP towards the base and redistributes blood from the LA to the LV via the mitral valve. When it reaches the switching threshold -S_D_, the contraction force *F_C_* starts to work in the opposite direction, which represents ventricular contraction. In this way, the AVPD leverages longitudinal pumping that results in ejection of blood to the Aorta. The ventricular contraction stops as soon as the AVP reaches the threshold *S_D_*. A relaxation phase follows where *F_C_* equals zero and the AVP moves slowly to its original position. This longitudinal pumping is well described by a piston unit concept, where the piston is placed between LA and LV with constant crosssections *A*_LA_ and *A*_LV_ respectively. We illustrate this piston representation at the bottom of Figure 2. The AVP model assumes that longitudinal pumping is supported by the radial squeezing of LV walls.

**Fig. 2.**
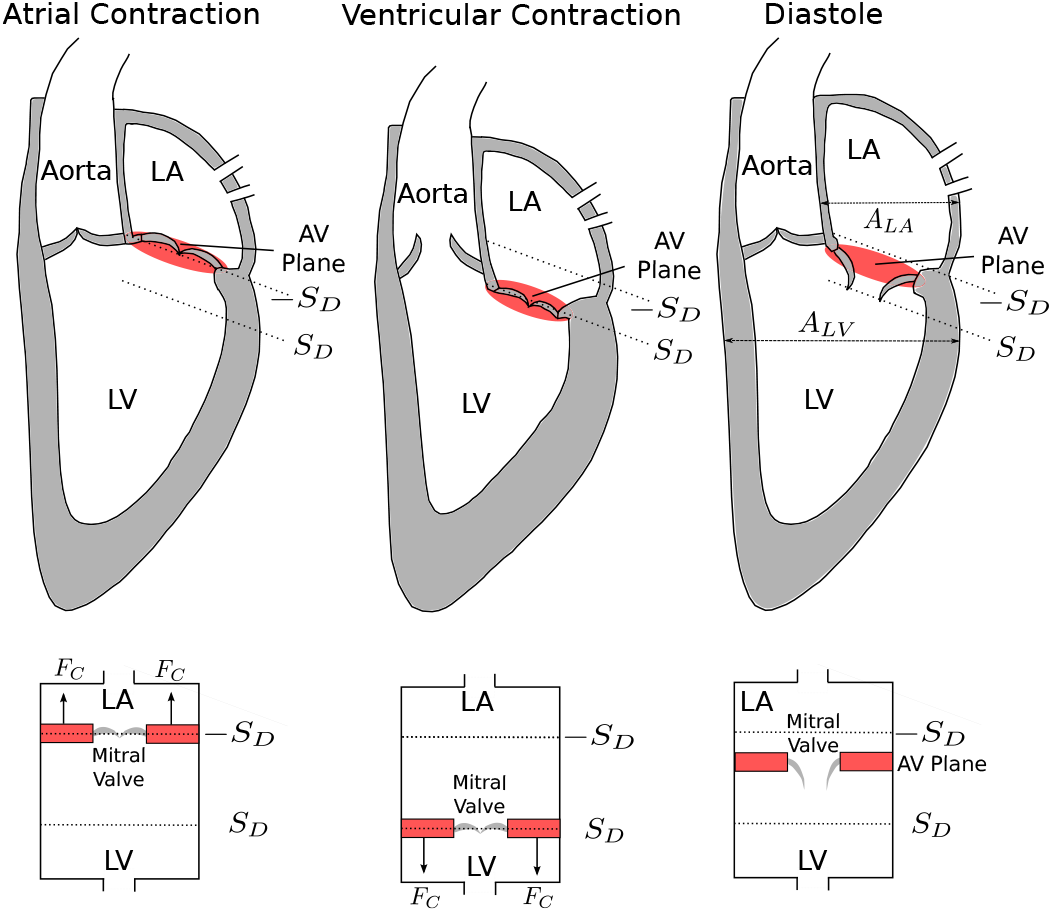
Illustration of the atrioventricular plane displacement concept for the left heart. The atrioventricular plane moves forth and back between —*S_D_* and *S_D_*, pulled by the contraction force *F_C_* resulting in blood redistribution from LA to the LV to the Aorta. This behavior resembles a piston pump, as depicted at the bottom, where —*S_D_* and *S_D_* mark the longitudinal displacement into basal and apical direction respectively.

### B. Physiological assumptions and constraints

This study makes a series of assumptions, which are explained here. We list the applied constraint parameter values in Supplemental Material II.

1. *Dilated left heart failure:* The proposed model is adapted in order to represent a typical LVAD patient candidate’s heart situation [9]. This includes modeling left-sided heart failure with decreased cardiac output and dilated cardiomyopathy^2^ with enlarged LA and LV. To this end, we modify certain model parameters, including an increased compliance and increased cross-sectional area of LV as described in more detail in Supplemental Material I. In addition, further parameters can be adapted to a specific patient, as explained in Chapter II-C.
2. *Steady-state situation:* We assume the cardiovascular and circulatory system is in *steady-state,* in the sense of

- there are no rapid or major changes of cardiac output and the heart cycle length,
- the system has already adapted to the LVAD implementation,
- the patient is at rest. These assumptions justify to neglect the autoregulatory mechanisms of cardiovascular pumping such as the systemic baroreflex feedback process and beat to beat myocardium wall strain adaptation based upon the Frank-Starling effect.
3. *Feasible instantaneous pump speed changes:* In practice, due to blood inertia, it is impossible to arbitrarily adjust the pump speed. Here we neglect blood and rotor inertia effects and assume that the pump speed can be varied without restrictions. Connected to this, blood is considered as Newtonian fluid and no blood rheology changes are taken into account.
4. *Blood inflow equals outflow:* In conjunction with the steady-state assumption, we require that the amount of accumulated incoming blood in the LV is equal to the accumulated amount of blood ejected out of the LV over the time horizon [*t*_0_, *t_f_*]. For this purpose we introduce the tolerance parameter *ϵ_flow_* > 0 and define (with *Q*_MV_(*t*),*Q*_Aov_(*t*) ≥ 0) the constraint:

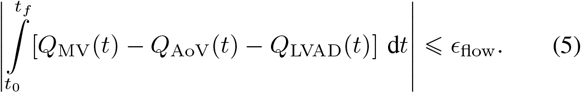
5. *Periodicity of the heart cycle:* The steady-state assumption implies that it is sufficient to consider only one heart cycle since there are no significant differences between several heart cycles. Thus, in this study we fix the time horizon to the length of one heart cycle. In this way, the steadystate condition translates into a periodicity constraint denoting that the differential state values at the beginning of the heart cycle should be equal to the ones at the end of the cycle. In mathematical terms this results with *ϵ*_per_ > 0 in

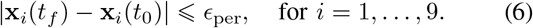
6. *Partial LVAD support:* When using an LVAD in the clinical setting, a distinction is made between full and partial support. While the LV does not contribute to blood ejection with full support, the aortic valve still opens with partial support because the LV contraction force is still strong enough to pump *partially*. We assume partial support, that is:

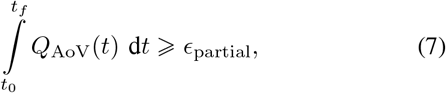

with *ϵ*_partial_ > 0.
7. *Back flow of blood from the Aorta in the LV:* We want to restrict the back flow from the Aorta in the LV via the LVAD. For this, we introduce the tolerance *ϵ*_back_ > 0 and require

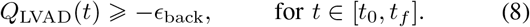
8. *Frank-Starling like control:* One objective of using an LVAD is to provide sufficient perfusion to the patient’s body. The Frank-Starling mechanism regulates physiologically the cardiac output, i.e., the amount of blood ejected by the LV into the Aorta per minute, according to the current need. We consider this mechanism as a desirable mode of operation and seek a pump speed control policy that results in an actual cardiac output that equals approximately a desired cardiac output *V*_CO_ ∈ ℝ_+_:

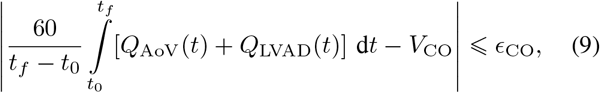

where *ϵ*_CO_ > 0. We note that the actual cardiac output should not exceed the desired cardiac output up to the tolerance, since this could result in fatigue for the patient.
9. *Variable bounds and suction prevention:* We require the differential state variables and the pump speed control to be in realistic ranges. Let x_lb_, **x**_ub_ ∈ ℝ^9^, respectively *u*_lb_,u_ub_ ∈ ℝ denote appropriate lower and upper bounds. The box constraints read

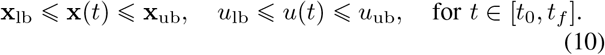 In this way, we are able to prevent the occurrence of suction, which describes the situation of excessive pumping that may cause a collapse of the ventricle if *P*_LV_(*t*) is very low.

### C. Clinical data and Model Personalization

This study uses data that were obtained retrospectively from the University Hospital Magdeburg, Department of Cardiology [50]. An exemplary subject was selected who involved a dilated LV and suffered from systolic left-sided heart failure. Data collection was performed via conductance catheterization for LV pressure measurements and via echocardiography for other data. The subject showed in rest a heart frequency of 67 beats per minute with a cardiac output of about 3.5 liters per minute. Further hemodynamic characteristics of the selected subject are shown in Table I.

**TABLE I.** Measured hemodynamic data for the example subject. Pulmonary capillary wedge (PCW) pressure represents a surrogate for LA pressure.

| Parameter | End-systolic | End-diastolic |
| --- | --- | --- |
| LV volume | 281 ml | 228 ml |
| LV pressure | 120 mmHg | 5 mmHg |
| PCW pressure | 28 mmHg | 14 mmHg |
| Aortic pressure | 121 mmHg | 53 mmHg |

We selected a representative cardiac cycle with the duration *h*_cycle_ = 0.89 seconds and 27 measured data points.

We propose to personalize the model via a parameter estimation (PE) method. For this, we formulate an optimization problem with the model equations as constraints and a nonlinear regression term as objective that minimizes the difference of model response values to the measured subject data. Here, we minimize the difference between measured LV pressure for selected time points and their corresponding model output values, however, this approach can also be applied to a general measured data set with more differential state types involved. We denote with 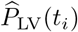 the measured LV pressure at time point *t_i_* ∈ [*t*_0_,*t_f_*]. We choose the parameters to be estimated as proposed in [27] with high sensitivity with respect to the LV pressure. These parameters are

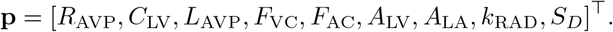

We bound the parameters to be in a realistic range, i.e., **p**_lb_ ≤ **P** ≤ **p**_ub_, see Supplemental Material II for further details. The selected subject had not (yet) implanted an LVAD, so we set *Q*_LVAD_(*t*) to zero and neglect the control *u*(*t*) and constraints on *Q*_LVAD_(*t*) for the PE. The parameter (point) estimation problem is defined as the following optimization problem:

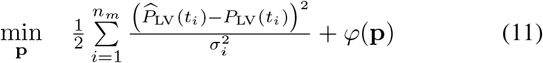

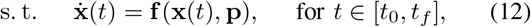

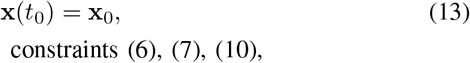

constraints (6), (7), (10),
where *n_m_* = 27 denotes the number of available measurements, **x**_0_ the initial values, and *σ_i_* is the standard deviation of the measurement at time *t_i_*, here set to one. The term *ϕ*(**p**) allows to incorporate a priori information of the parameters, which we here set to zero^3^.

### D. Optimal Control Problem Formulation

Based on a personalized model, we take interest in an advantageous application of the LVAD for a (possible) patient. An OCP offers the framework to include generic constraints and objective functions. While we have already defined the constraints in Section II-B, for the objective we reuse the multiobjective function from [13]. This objective constitutes a compromise function that aims for ventricular unloading and ensures the opening of the aortic valve. A permanent closure of the aortic valve may lead to fusion of the aortic valvular cusps and a resulting thrombus formation [10]. By ventricular unloading we refer to reducing the hydraulic work that the LV has to perform in order to provide sufficient perfusion. Let *ϱ*_1_ ∈ [0,1] denote a weighting parameter that facilitates to put one objective more into focus and let *ϱ*_2_ and *ϱ*_3_ denote unit scaling factors, see Supplemental Material II for more details. Then, we introduce the objective as

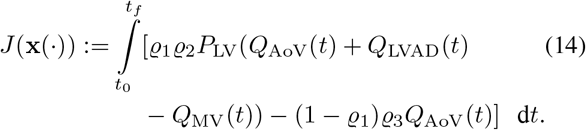

The first term accounts for the ventricular unloading, while the second term causes aortic valve opening via maximizing the flow through this valve. We consider the following optimization problem, where we minimize the above objective over the differential states **x**(·) and the continuous control *u*(·):

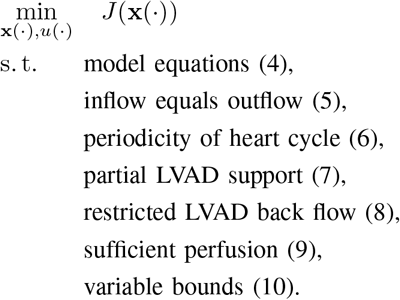

For this optimization problem we investigate three different scenarios regarding the pump speed control.

1. *Constant speed:* This represents the usual clinical setting and is expressed by *u*(*t*): = *u*_con_ ∈ [*u*_lb_, *u*_ub_] for *t* ∈ [*t*_0_,*t_f_*].
2. *Continuous speed:* There are no restrictions on *u(·)* apart from lower and upper bounds.
3. *Pwc speed:* In order to create pulsatility, this scenario considers to switch between different constant speed modes. For this we use the indicator function notation

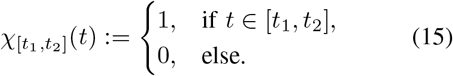 We assume *u*(·) to be a step function with three different levels *u*_1_,*u*_2_,*u*_3_ ∈ [*u*_lb_,*u*_ub_]:

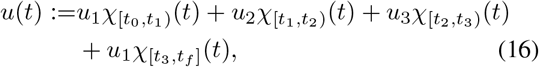

where *t*_1_, *t*_2_, *t*_3_ are switching times to be determined^4^. We require minimal time durations for the different speed levels because rapid changes are not feasible in a realistic setting. Let *D*_1_, *D*_2_, *D*_3_ > 0 denote these so-called minimum dwell times and we introduce the constraints

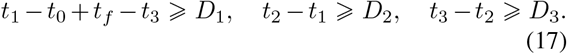

### E. Algorithmic Approach

We distinguish between *explicit* and *implicit* switches that result in discontinuous variables for the PE and the OCP. The pump speed control should be in one scenario pwc; however, since we can control when this switch occurs between different speed modes, we call this switch *explicit*. By *implicit* switches we refer to changes of the model equations in **f** that happen as soon as the differential states satisfy certain conditions. The valve flows and the contraction force induce such implicit switches. While the valve switches are defined in (2a)–(2b), we specify the contraction force switches in the following.

#### 1) Implicit switches through the contraction force

As we consider only one heart cycle, the atrial and ventricular contraction takes place once. We assume a physiological order, that is atrial before ventricular contraction followed by a relaxation phase. Initially, let -*S_D_* ≤ *s*(*t*_0_) ≤ *S_D_*. We further assume the following switching times exist:

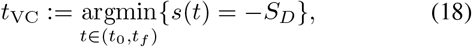

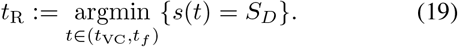

Then, the contraction force is defined as

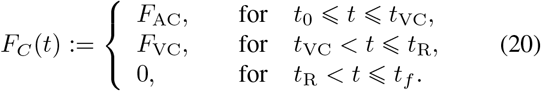

#### 2) Dividing the cardiac cycle into phases

The periodic switching nature of the cardiac cycle model makes the solving process challenging. We need to identify when switching happens and what the successive active subsystems of **f** are. If we combine all possible valve positions and contraction force settings, we get 12 different subsystems. To reduce complexity we assume a specific sequence of active subsystems for the cardiac cycle taking advantage of physiological relationships in the human heart. Thus, we divide the heart cycle into seven phases similar as in [27]. Table II and Fig. 3 explain the phases of the ordered sequence, where the switching times are denoted with *τ_i_, i* = 1,…, 6.

**Fig. 3.**
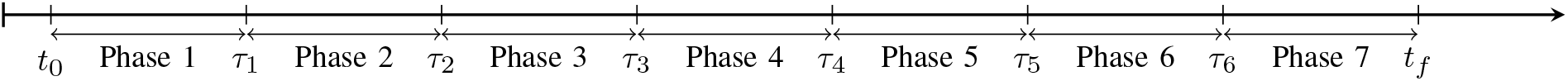
Time course of the assumed active phase sequence with switching events. The switching times *τ_i_* are variables in the optimization problem.

**Table II.** Assumed sequence of active phases. For instance, in the first phase the LA contracts, the mitral valve is open and the aortic valve is closed.

| Phase | $F_C$ mode | Mitral valve | Aortic valve |
| --- | --- | --- | --- |
| 1 | AC | open | closed |
| 2 | VC | open | closed |
| 3 | VC | closed | closed |
| 4 | VC | closed | open |
| 5 | 0 | closed | open |
| 6 | 0 | closed | closed |
| 7 | 0 | open | closed |

The modes from Table II translate into the following con-straints for the optimization problem and for **f**:

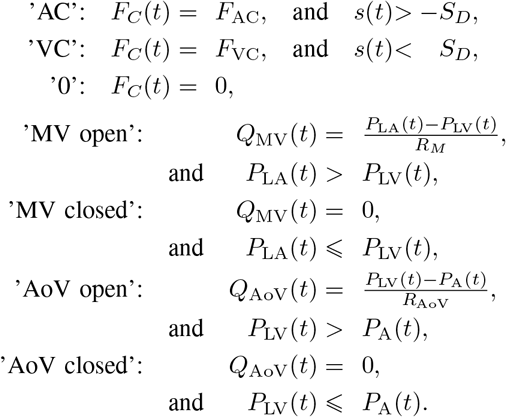

By fixing the sequence of active subsystems, the PE and OCP transform into multiphase problems [51], where only the switching times needs to be determined.

#### 3) Switching time optimization

We use switching time optimization [51], [52] to determine the switching times *τ*_1_,…,*τ*_6_ so that we can transform the originally discrete optimization problems into continuous ones. The idea of switching time optimization relies on a time transformation *t* = (*t*_2_ — *t*_1_)*τ*, which exploits that

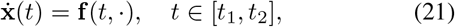

is equivalent to

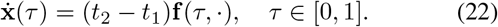

With this idea, we reformulate the multiphase model equation:

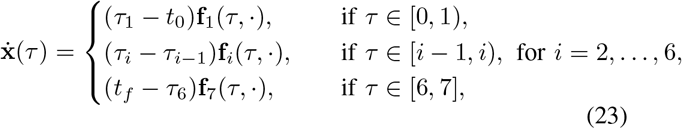

where **f**_*i*_ denotes the model equation for the *i*th phase. We notice that the phase durations enter the equation via (*τ_i_* – *τ*_*i*-1_) as continuous variables. At the end of each phase, the switching constraints for the contraction force and the valve flows as described in Section II-E.2 need to be fulfilled at the transformed switching time points up to a tolerance *ϵ*_sw_ > 0:

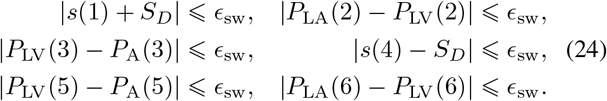

This switching time reformulation provides the framework to solve the PE problem and the OCP with constant and continuous pump speed as optimization problem with solely continuous variables. For the pwc pump speed modulation we also need to find the switching times *t*_1_,*t*_2_,*t*_3_ between the three different speed levels as introduced in Section II-D. Here, we assume

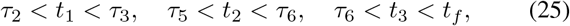

i.e., the first speed change occurs between mitral valve closing and aortic valve opening, the second between aortic valve closing and mitral valve opening, and the third between mitral valve opening and the end of the heart cycle. Thus, we divide the third, sixth and seventh phase from Section II-E.2 into two phases each so that in total nine switching times for ten phases need to be determined.

#### 4) Numerical solution of optimization problems

We use direct collocation [53] to transform the continuous time optimization problems via temporal discretization into nonlinear programs (NLPs). We apply an equidistant discretization grid with grid length of 1 ms and we let the control values change their values only on the grid points. The differential state trajectories are approximated with Radau collocation polynomials [54] of degree 3. We implemented the optimization problems in `python` v3.7.5 and used `CasADi` v3.4.5 [55] to parse the resulting NLP with efficient derivative calculation of Jacobians and Hessians to the solver `IPOPT` v3.12.3 [56]. For the PE problem we applied the Gauss-Newton method [54] so that calculation of Hessians is not required.

The lengths of the model phases were extracted from pressure time series and other continuous data and used for initialization of the switching times *τ_i_*. Theses phase durations were fixed for the PE problem and set variable for the OCP. We further initialized the PE problem with variable values based on a simulation with default parameter values, see Supplemental Material I. The OCP was initialized with simulated variable values obtained with estimated parameters and constant pump speed equal to 8000 rpm.

## III. Numerical Results

### A. Patient specification

Solving the PE problem from Section II-C resulted in the values

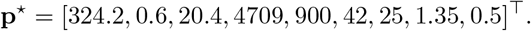

The result of the switching distance parameter, i.e. *S_D_* = 0.5, is equivalent to an AVPD of only 10 mm and thus indicates a reduced ventricular function. The situation of heart failure is reflected well by the estimated parameter values. Particularly, the LV compliance is increased, the amplitude of the contraction forces *F*_AC_ and *F*_VC_ is decreased and the parameter *k*_RAD_ accounting for relative contribution of radial pumping is increased. Fig. 4 shows the measured data points 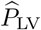 together with the obtained *P*_LV_ from the PE solution.

**Fig. 4.**
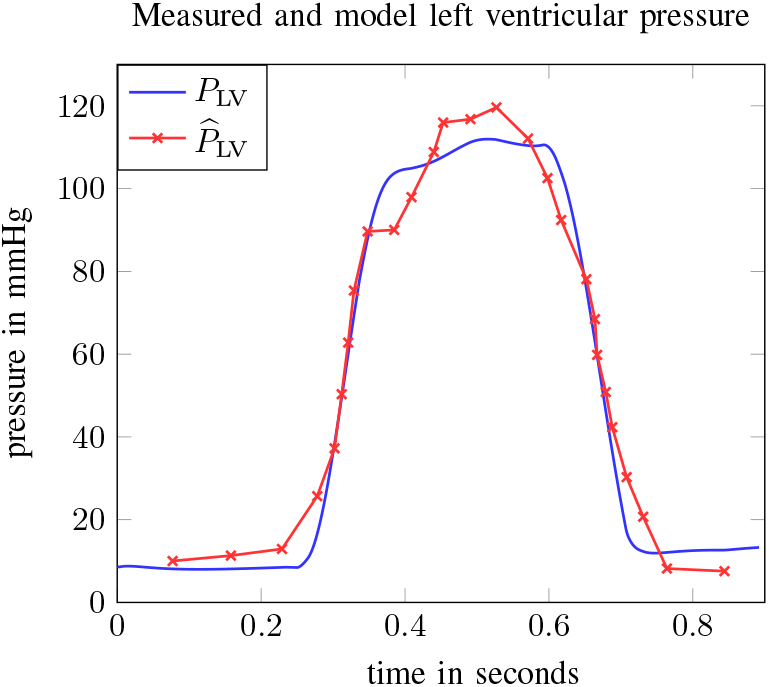
Measured 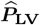 values and resulting *P*_LV_ trajectory based on the parameters obtained from solving the PE problem from Section II-C.

The model response *P*_LV_ reflects the measured data points especially with respect to the duration of ventricular contraction, while its peak is slightly underestimated. The transcripted nonlinear regression problem was solved by `IPOPT` after 230 seconds and with an objective value of 512 mmHg^2^, which is equivalent to a root-mean-square deviation of 6.16 mmHg. We observed numerical instabilities when solving the PE problem. The convergence of the algorithm seems to depend heavily on the initial solution, which stresses the importance of the proposed initialization from Section II-E.4. In addition, we have used mild termination criteria for `IPOPT` and chose an big tolerance value for the periodicity constraint (6). Fig. 5 depicts all model pressure trajectories based on the PE and the six switching times between the different model phases. We observe that the aortic pressure *P_A_* resembles the LV pressure *P*_LV_ during ventricular systole and the systemic pressure *P*_S_ else, apart from some small oscillations. Likewise, *P*_LA_ and *P*_V_ represent similarly high pressures, although *P*_LA_ adopts to *P*_LV_ depending on the mitral valve opening.

**Fig. 5.**
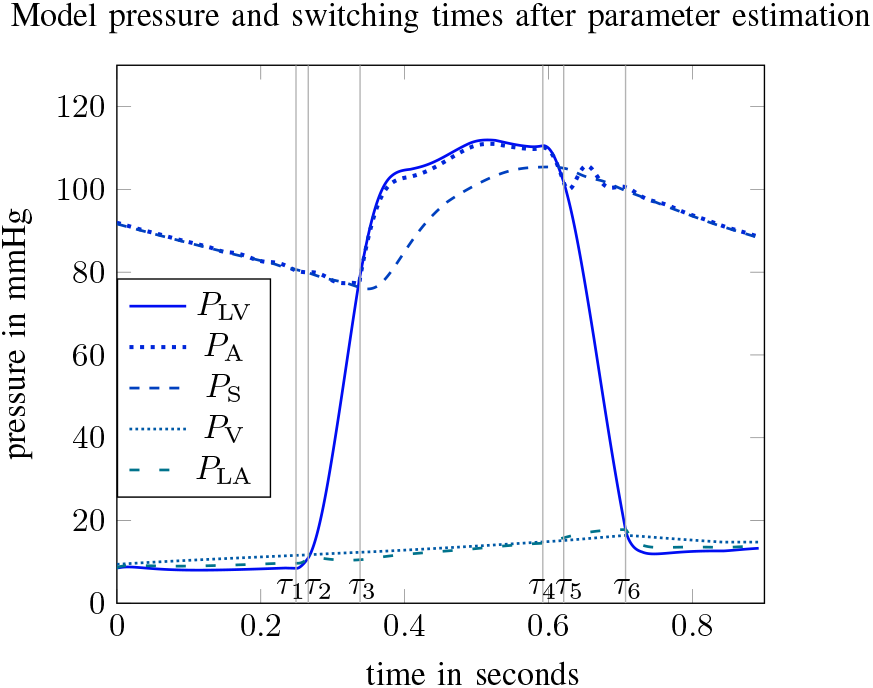
Simulated pressure functions based on the parameters obtained from solving the PE problem from Section II-C. The switching times *τ_i_* are depicted with the vertical grey lines.

### B. Pump control policies

We solved the OCP according to the proposed algorithm from Section II-E. The applied parameters for the objective and constraints such as the tolerance parameters *ϵ*., the dwell times *D*., and lower and upper bounds on variables are listed in Supplemental Material II. Fig. 6 shows the pressure functions for the three pump speed scenarios. The outcomes of the continuous and pwc scenario are very similar. They show an elevated peak of *P*_LV_ compared to the parameter estimated solution from Fig. 5. While the rise and fall of the pressure profiles before and after the ventricular contraction is significantly steeper than with the parameter estimated solution, its duration, i.e. *τ*_4_ — *τ*_1_, is in a similar range due to an enforced minimum dwell time of 0.2 seconds for the ventricular contraction. Very low values occur for *P*_LV_ directly before *τ*_1_, however, they are still above the threshold for suction. We notice that the cycle duration for the pwc scenario is 0.88 seconds and slightly longer than for the other two scenarios with a duration of about 0.84 seconds, as we allow a slight deviation of 50 ms from the standard cycle length. The constant speed scenario involves also an increased aortic and ventricular pressure compared to the parameter estimated solution, though their peaks are significantly lower compared with the continuous and pwc speed scenarios.

**Fig. 6.**
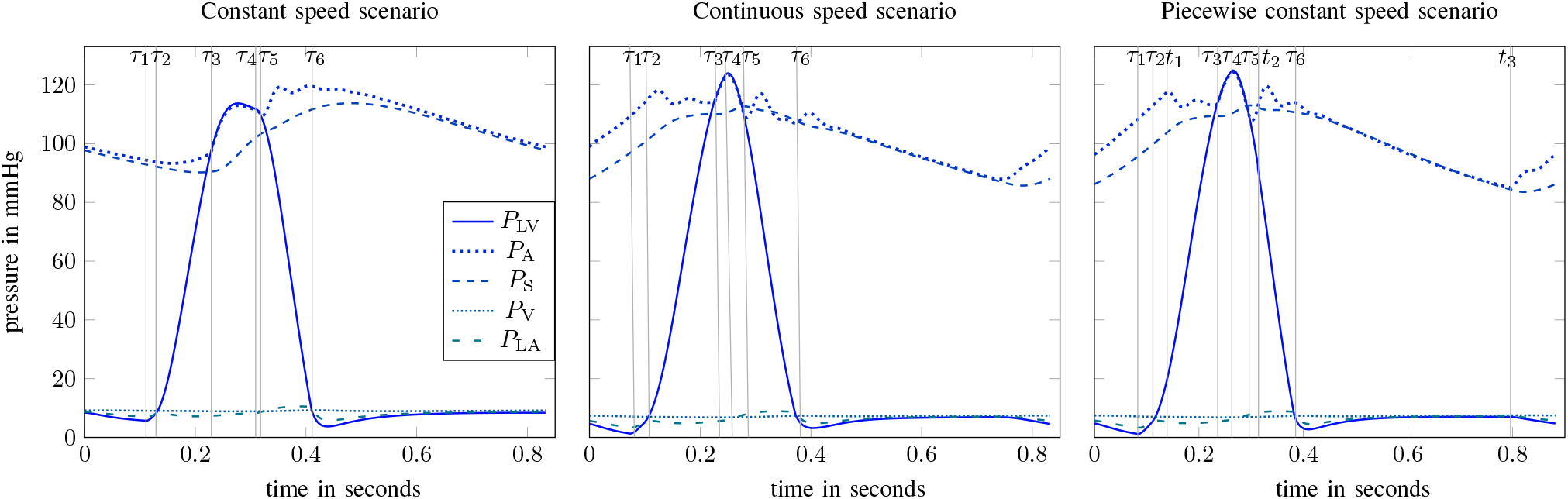
Calculated pressure differential states for the OCP solutions with different pump speed scenarios. The switching times *τ_i_* and pwc speed changing times *τ_i_* are depicted with the vertical grey lines.

Fig. 7 illustrates the different optimal pump speed profiles and the according results for the flows, AVPD speed, and AVPD position. We observe that the continuous speed profile provides counterpulsative pump support and the pwc speed profile approximates this profile. Due to this similar pump speed behavior, the optimal differential state trajectories resemble each other. The main portion of the flows through the LVAD and the aorta appears for constant pump speed during ventricular contraction. In contrast, with continuous or pwc speed, large flow values occur already during atrial contraction followed by a peak during ventricular contraction, which accounts for the remaining physiological contraction force. The flow for the continuous pump speed appears to be slightly negative around *t* = 0.4 since we relaxed the tolerance parameter in (8) for achieving numerical convergence. The upper right panel shows that the larger amplitudes of the pumping speed for the continuous and pwc scenario compared with constant speed translate into faster AVP movements.

**Fig. 7.**
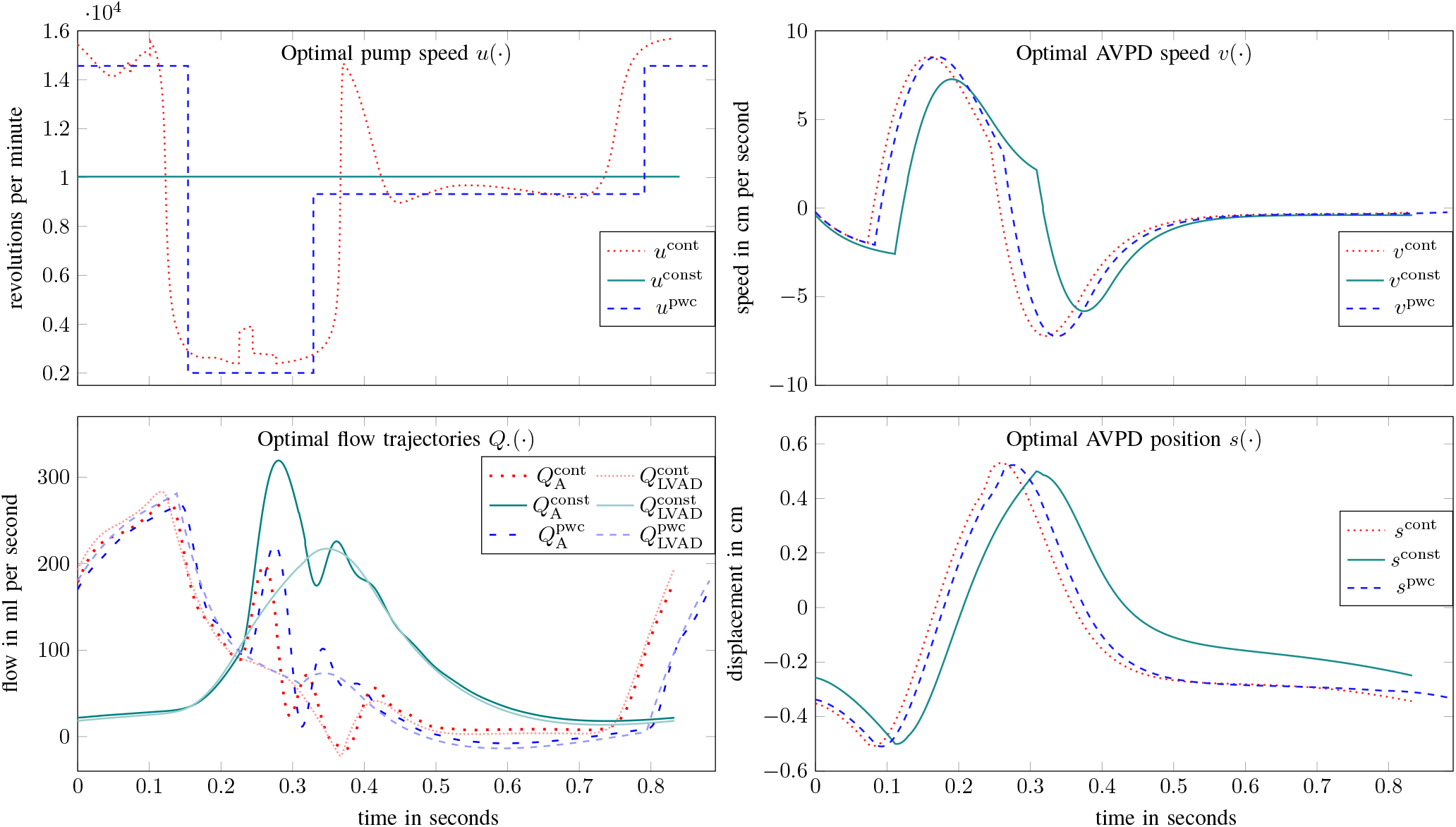
Calculated pump speed and differential states for the OCP solutions. The superscripts *cont, const*, and *pwc* abbreviate *continuous, constant* and *piecewise constant* rotor speed.

Fig. 8 summarizes the objective values and runtimes for the OCP solutions. Clearly, the obtained objective value with the pwc speed profile is only slightly larger than the one calculated with continuous pump speed, while the objective value with constant speed is not competitive.

**Fig. 8.**
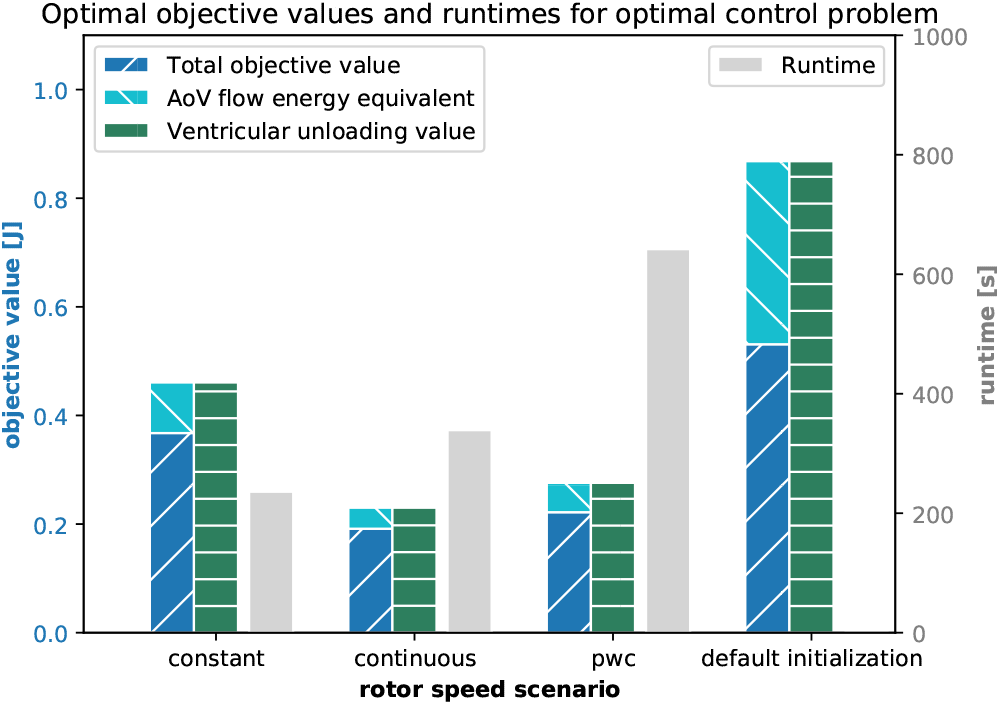
Comparison of constructed optimal objective values and runtimes for the OCP from Section II-D. The objective values for the default initialization are obtained with *u*(·) ≡ 8000rpm. Notice that the total objective value results from subtracting the aortic valve flow value from the ventricular unloading, as defined in (14). The objective value decreases from 0.53105 after initialization to 0.36758 with constant speed. Continuous and pwc speed modulation construct even lower objective values, which amount to 0.19149 and 0.22182 respectively.

## IV. Discussion

The root-mean-square error between 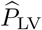 and *P*_LV_ is about twice as large as the one constructed via a trial method in [27]. However, the latter study did not take into account the constraints (6), (7), and (10) during the data fitting procedure. The performed PE should be discussed critically with respect to overfitting since we optimized nine parameters with available measurements for only one differential state. To this end, we calculated the relative standard deviation (SD) values based on the Fisher-Information matrix as defined in [57], which represents a common criteria for evaluating the quality of the results of a PE. The SD values with respect to the optimal parameter vector **p**\* are

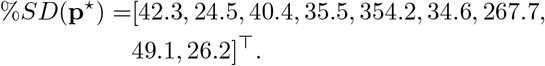

The calculated SD values are mostly quite low and thus indicate a robust quality of the obtained estimation. In contrast, the values for *F*_AC_ and *A*_LA_ with about 354 % and 268 % are quite large and we postulate that they have a minor impact on *P*_LV_. Future work should focus on sensitivity analysis for choosing the right parameters to be estimated.

The increased peak of the LV pressure function after using pumping assist is consistent with typical partial LVAD support [18]. The causality can be explained as follows: The LVAD delivers continuously blood to the aorta, increasing the aortic pressure. Therefore the LV pressure must be elevated for the aortic valve to open. This effect is more pronounced in the scenarios with continuous and pwc pump speed, which can be categorized as counterpulsative policies and thus facilitate the aortic pressure to oscillate less. Our finding that a counterpulsative pumping strategy minimizes the given objective function is in accordance with [39], where the same objective function was applied.

After applying the LVAD support, we observed shortened atrial contraction phases, which is equivalent to shortened ventricular filling phases. We interpret this behavior as a result of maximizing the ventricular unloading. This implies the LA pumps against less resistance and reaches the maximum contraction state more quickly, which is represented by the AVP reaching the switching distance —*S_D_*. In this way, the AVPD model can realistically capture interactions of an LVAD and the cardiac system, as elaborated in [42], [43]. The continuous and pwc pump speed scenarios have more degrees of freedom compared with the constant speed scenario and thus improve the diastolic function even more as depicted by the shortened atrial contraction phases.

### A. Connection to clinical application

LVADs work in an online environment where the system state can change rapidly, especially the heart rate and blood volume shift. Currently, there are no sensors available that provide long-term measure signals of the presented differential states [19]. Despite these aspects and our restricting assumptions (see next subsection), we claim that optimized speed profiles from offline computations may provide superior performance (see Fig. 8) and could be considered in the following way.

1. The optimal control framework can be used to benchmark a whole range of speed profiles, in particular modern pwc speed profiles, that result from different objective functions, models, and constraints.
2. These evaluations can be carried out on a patient-specific basis. As a proof of concept, we demonstrated for one patient that the cardiovascular model can be efficiently altered to represent the patient’s LV pressure function. This approach can be extended to include time series measurements of the aortic and LA pressure (via PCW pressure and conductance catheterization), the flows at the mitral and aortic valve, and the AVP speed and displacement (all via Doppler echocardiography). Overall, at least five differential states could be used for model personalization. The data used so far were measured invasively. In contrast, in routine clinical investigations, echocardiography can be used to measure and use non-invasive data concerning approximated PV loops and time series of valve flows.
3. Offline computations of an optimal speed profile for different situations, e.g. rest, exercise or rhythm disturbances such as atrial fibrillation, could be done beforehand and used in an online setting assuming information about the system status is available.
4. The presented algorithmic idea can be extended to model predictive control. In particular, the possibility to incorporate minimum dwell time constraints to avoid rapid speed changes paves the way for a realistic extension to the online setting.

### B. Limitations

#### 1) Simplified Model

We applied a lumped model that simplifies the heart and the cardiovascular system by neglecting the right heart, the pulmonary system, valve regurgitation and spatial interactions between compartments. Dilated heart failure is very commonly associated with valve regurgitation so that our assumption to neglect it should be seen critical. The passive LV compliance *C*_LV_ is set to be constant, but may change according to the Frank-Starling law. The way we model the LVAD and its interaction with the heart is also highly simplified. Moreover, neurological feedback processes such as the baroreflex are not captured.

#### 2) Assumptions

In reality, the heart rate and thus the duration of the cycle is very variable, especially through exertion or sport. In most cases of patients requiring the VAD therapy the stable heart rhythm is a scarce phenomenon. The rhythm disturbances are rather the dominant pattern in the individuals suffering from heart failure. Therefore the steady state assumption must be viewed critically. Furthermore, we neglect rotor and blood inertia so that the rotor speed can be controlled arbitrarily. Nevertheless, our framework may be good as a general starting point for the twofold development as part of future work. First we may be able to base the optimization process on critical parameters with respect to clinical availability. Second the early detection of atrial fibrillation as the most common rhythm disturbance in heart failure could be implemented to switch the working regime of the pump into different mode [58].

#### 3) Measured Data

We included in this study only one subject and measured data for only one differential state. Future work shall address numerical tests with several patients and with additional measured states.

#### 4) Control approach

Other measures of ventricular work such as the PV area could be applied for the objective function. This study assumes a canonical order of the active phases with respect to the valves and the contraction force. This order might not always be true in practice so that future work should consider optimization with implicit switches, but without fixing the order of active phases. Besides, we optimize over one heart cycle, whereas the pulsatility speed mode profiles of some LVADs such as HeartMate 3 last for more than one heart cycle.

### C. Switched Systems framework applicability

The developed multiphase algorithm is applicable to other models and settings. For example, the OCP can also be interpreted on a cardiac model with time-varying elastance function as a switched system, with the valves still representing the implicit switches and changes of the constant pump speed representing “controllable” switches. Analogously, the framework can be beneficial for PE of cardiac models without LVAD application, but with different scope, e.g. cardiac resynchronization therapy.

There are similar devices to an LVAD available or under development for which an OCP could be solved efficiently with the switched systems framework. For instance, total artificial hearts such as RealHeart [59] and Carmat [60] or intra-aortic blood pumps [61] involve also discrete system changes, induced by piston pumps (RealHeart), controlled valves (Carmat) or pulsatility rotor speed modes (intra-aortic blood pump). Finally, the next generation LVADs may include more advanced control features that can lead to different control modes to switch on/off. The TORVAD device [62] falls into this category and works with two magnetic pistons within a torus generating pulsatile flow.

## V. Conclusion

We have proposed a novel switched systems algorithm for the optimal control of LVADs that provides the opportunity to calculate optimal constant, pwc, or continuous pump speed profiles. As a proof of concept, we showed that this algorithm can be used to adapt a cardiovascular model to patient specific data and to benchmark simulations of personalized LVAD control policies. The importance of achieving hemodynamic optimization in LVAD patients is highlighted by a significantly lower rate of hospital readmissions [12], and could benefit from in silico analysis such as the presented speed profile evaluations. Moreover, we have demonstrated realistic simulations of a model that is based on AVPD instead of using the widespread time-varying elastance model and examined thereby the heart to LVAD interactions. Future work may test the algorithm on more patient data, more realistic conditions such as exercise or rhythm disturbance, and with model extensions. The proposed algorithm could be also beneficial for the evaluation of pulsatile speed modulation modes of modern devices such as HeartMate 3 or HeartWare HVAD.

## Acknowledgment

The authors thank Felix Gnettner for fruitful discussions on modeling cardiovascular system dynamics. This project has received funding from the European Research Council (ERC, grant agreement No 647573) under the European Union’s Horizon 2020 research and innovation program and and from Deutsche Forschungsgemeinschaft (DFG, German Research Foundation) – 314838170, GRK 2297 MathCoRe and SPP 1962.

## Supplemental Material

### I. Model parameter values

The resistance and inertance of the LVAD is captured by the parameters *R_LVAD_* and *L_LVAD_*. We use the values from [49] as shown in Table Table S1 and given by

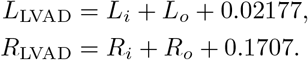

We altered the model from [27] to the situation of left heart failure by increasing the left atrial and ventricular compliance, increasing the systemic resistance, decreasing the length of AVPD, increasing the cross-section of LA and LV in order to account for a dilated heart, and decreasing the contraction force *F_C_*. The specific parameter values are defined in Table Table S1.

**TABLE Table S1.** Default parameter values for the cardiovascular, circulatory system, and LVAD model.

| Parameter | Description | Value |
| --- | --- | --- |
| $C_{LA}$ | Left atrial compliance | 0.6 ml/mmHg |
| $C_{LV}$ | Left ventricular compliance | 0.45 ml/mmHg |
| $C_A$ | Aortic compliance | 0.08 ml/mmHg |
| $C_S$ | Systemic arterial compliance | 1.2 ml/mmHg |
| $C_V$ | Venous compliance | 50 ml/mmHg |
| $R_M$ | Mitral valve resistance | 0.005 mmHg s/ml |
| $R_{AoV}$ | Aortic valve resistance | 0.005 mmHg s/ml |
| $R_S$ | Systemic arterial resistance | 1.1 mmHg s/ml |
| $R_C$ | Characteristic resistance | 0.05 mmHg s/ml |
| $R_i$ | LVAD inlet resistance | 0.0677 mmHg s/ml |
| $R_o$ | LVAD outlet resistance | 0.0677 mmHg s/ml |
| $R_{AVP}$ | Damping of AVP | 300 mmHg cm s |
| $L_S$ | Arterial inertance | 0.001 mmHg s<sup>2</sup>/ml |
| $L_i$ | Inlet inertance of LVAD | 0.0127 mmHg s<sup>2</sup>/ml |
| $L_o$ | Outlet inertance of LVAD | 0.0127 mmHg s<sup>2</sup>/ml |
| $L_{AVP}$ | Inertia of AVP | 30 mmHg cm s<sup>2</sup> |
| $\beta$ | Pump-to-pressure coefficient | -9.9025e-7 |
| $A_{LA}$ | Left atrial cross-section | 25 cm<sup>2</sup> |
| $A_{LV}$ | Left ventricular cross-section | 50 cm<sup>2</sup> |
| $F_{AC}$ | Left atrial contraction force | 1000 mmHg cm<sup>2</sup> |
| $F_{VC}$ | LV contraction force | 5000 mmHg cm<sup>2</sup> |
| $k_{RAD}$ | Radial function coefficient | 1.2 |
| $S_D$ | Switching threshold for AVP | 0.4 cm |

## II. Optimization parameter values

The following lower and upper bounds for the variables of the optimization problems were applied.

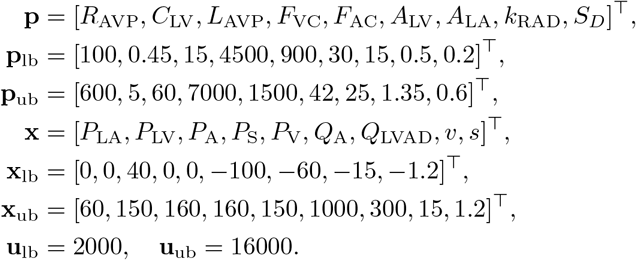

We used as initial differential state value for the parameter estimation problem:

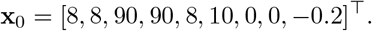

Furthermore, the following parameters for constraints were applied:

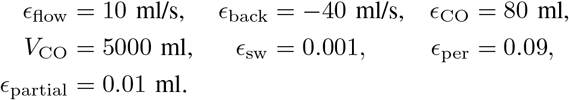

We permitted a deviation of 50 ms for the model heart cycle from the subject’s measured cycle length. The minimum dwell times for the pwc pump speed levels were

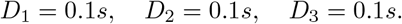

We also enforced upper bounds *τ*_ub_ on the seven phase durations to be

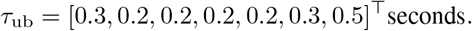

The objective parameters are

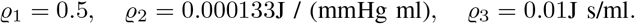

We used the following `IPOPT` setting: *‘tol’: 1e-6, ‘constr_viol_to?: 1e-6, ‘compl_inf_tol’: 1e-6, ‘dual_inf_tol’: 1e-6.*

## Footnotes

1 We neglect valve regurgitation and set the back flow to zero. We discuss this assumption in Section IV-B.

2 Alternatively, myocardial infarction is a common case related to LVAD patients; however, it is challenging to represent scars adequately with a zero-dimensional lumped model.

3 Future work should consider a priori information in the form of 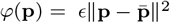 instead of imposing lower and upper bounds on **p**.

4 In fact, we can drop *χ*_[*t*_3_,*t_f_*]_(*t*) since **x**(0) is free. However, the algorithmic idea in the next section exploits a fixed sequence of active system phases so that we keep this term.

